# The neonatal Fc receptor and DPP4 are human astrovirus receptors

**DOI:** 10.1101/2024.07.12.603331

**Authors:** Harshad Ingle, Jerome M. Molleston, Paige D. Hall, Duyen Bui, Leran Wang, Lynne Foster, Avan Antia, Siyuan Ding, Sanghyun Lee, Daved H. Fremont, Megan T. Baldridge

**Affiliations:** Division of Infectious Diseases, Department of Medicine, Edison Family Center for Genome Sciences & Systems Biology, Washington University School of Medicine; St. Louis, Missouri, USA; Department of Pediatrics, Washington University School of Medicine; St. Louis, MO, USA; Department of Pathology and Immunology, Washington University School of Medicine; St. Louis, MO, USA; Department of Molecular Microbiology, Washington University School of Medicine; St. Louis, Missouri, USA; Department of Molecular Microbiology and Immunology, Division of Biology and Medicine, Brown University; Providence, RI, USA; Department of Biochemistry and Molecular Biophysics, Washington University School of Medicine; St. Louis, MO 63110, USA

**Author notes:** These authors contributed equally (Harshad Ingle, Jerome M. Molleston, Paige D. Hall).

## Abstract

Human astroviruses (HAstV) are major global causes of gastroenteritis, but little is known about host factors required for their cellular entry. Here, we utilized complementary CRISPR-Cas9-based knockout and activation screening approaches and identified neonatal Fc receptor (FcRn) and dipeptidyl-peptidase IV (DPP4) as entry factors for HAstV infection of human intestinal epithelial cells. Disruption of FcRn or DPP4 reduced HAstV infection in permissive cells and, reciprocally, overexpression of these factors in non-permissive cells was sufficient to promote infection. We observed direct binding between FcRn and HAstV virions as well as purified spike protein. Finally, inhibitors for DPP4 and FcRn currently in clinical use prevent HAstV infection in cell lines and primary human enteroids. Thus, our results reveal mechanisms of HAstV entry as well as druggable targets.

**One-Sentence Summary:** Targeting FcRn or DPP4 using available therapies effectively prevents human astrovirus infection in human enteroid cultures.

## Main Text

Human astroviruses (HAstVs) are major causes of pediatric gastroenteritis worldwide, with up to 10% of sporadic cases of non-bacterial diarrhea in children attributed to classical HAstVs^1–3^. There is currently no vaccine or approved antiviral treatment for these positive-sense, single-stranded RNA viruses, in part because little is currently known about the mechanisms of HAstV entry. Previous antigenicity and sequence analyses have suggested that the spike P2 domain of the viral capsid is likely to be involved in cell attachment and receptor binding^4–6^, but the host factors involved in this process have not yet been identified.

### FcRn and DPP4 are required for HAstV infection

To discover proviral factors required for HAstV infection, we leveraged pooled genome-wide CRISPR-Cas9 screening. Since HAstVs can be readily cultivated in Caco2 cells, an immortalized colonic adenocarcinoma cell line^7–9^, we introduced the human Brunello library^10^ into Cas9-expressing Caco2 cells. We infected this library with HAstV1 at multiplicity of infection (MOI) of 10 for 24 hours. Uninfected cells were sorted using a monoclonal anti-HAstV capsid antibody^11–13^ (**Supp Fig. 1A**). After genomic DNA extraction, sgRNAs were Illumina-sequenced and the resulting sequencing data was analyzed using MAGeCK^14^. The top hits from the genome-wide knockout screen for HAstV1 were Fc gamma receptor and transporter (*FCGRT*), dipeptidyl-peptidase IV (*DPP4*), beta-2-microglobulin (*B2M*), muscleblind-like splicing regulator 1 (*MBNL1*), and solute carrier family 35 member C1 (*SLC35C1*) (**Fig. 1A**).

**Fig. 1.**
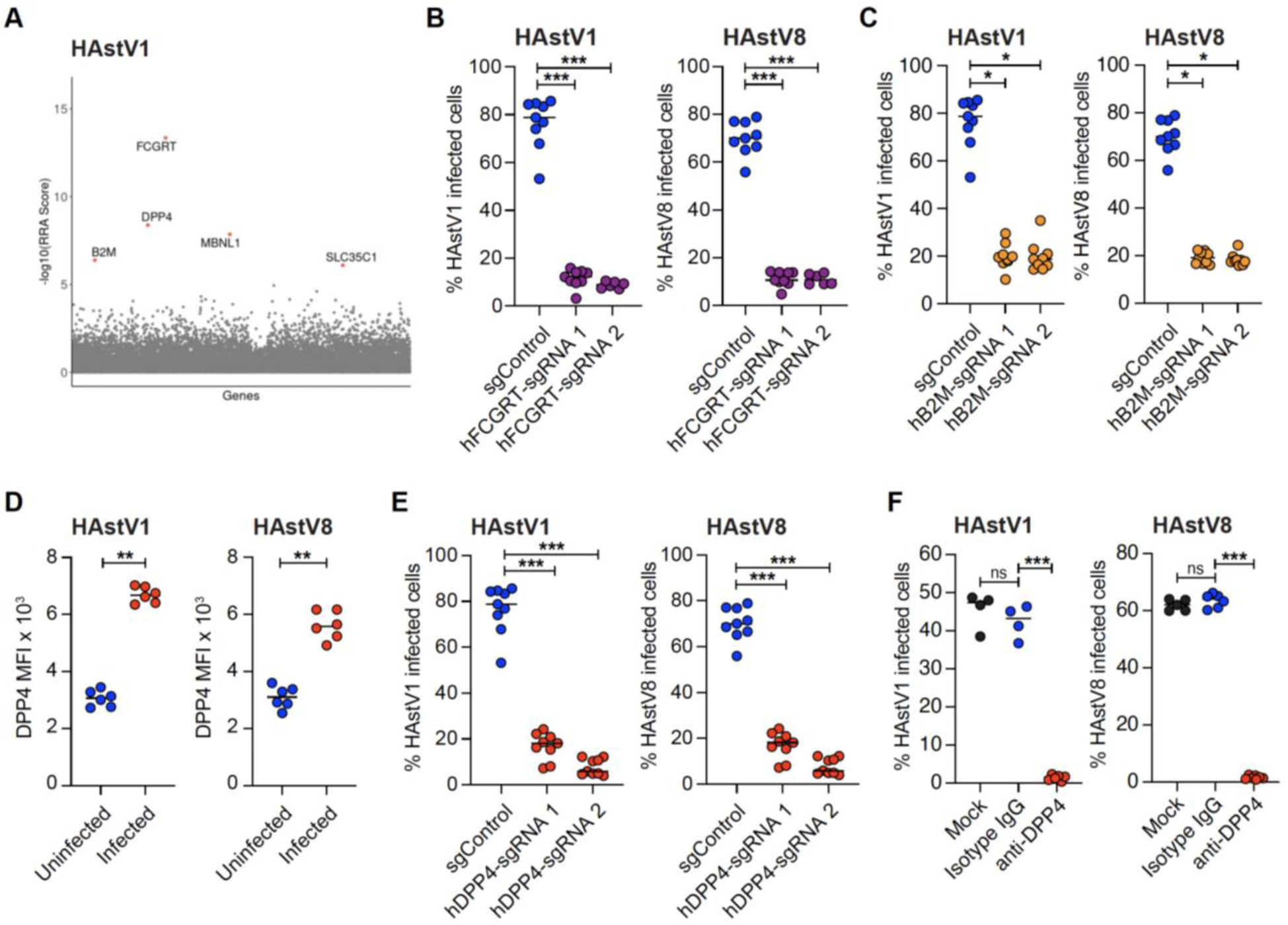
FcRn and DPP4 are required for HAstV entry and infection. (**A**) Enrichment [-log10 Robust Rank Aggregation (RRA)] scores of positively-selected sgRNAs in HAstV1-negative cells sorted 24hpi of a genome-wide CRISPR-Cas9 Caco2 cell library compared to HAstV1-positive cells, calculated by MAGeCK. (**B,C**) Cas9-Caco2 cells disrupted for *FCGRT* (n=6-9) or *B2M* (n=9) using two independent sgRNAs per gene or targeted with a control anti-GFP sgRNA (n=9) were infected with HAstV1or HAstV8, then stained with anti-HAstV capsid antibody at 24hpi. (**D**) Mean fluorescent intensity (MFI) of DPP4 in negative and positive cell populations of Caco2 cells infected with HAstV1 (n=6) or HAstV8 (n=6) infected at 24hpi. (**E**) Cas9-Caco2 cells were disrupted for *DPP4* (n=9) using two independent sgRNAs per gene or targeted with a control anti-GFP sgRNA (n=9), then assessed for infection using anti-HAstV capsid antibody 24hpi with HAstV1 or HAstV8. **(F**) Percentage of anti-HAstV capsid antibody-stained Caco2 cells treated with PBS (n=4-5) or isotype control (n=4-6) or anti-DPP4 polyclonal antibody (n=6-7) for 12h hours prior to infection with HAstV1 or HAstV8. Results from three independent experiments were analyzed using the Kruskal-Wallis test with Dunn’s post-test (B to F). *P<0.05; **P<0.01; ***P<0.001; ****P<0.0001. ns=not significant. Bars indicate mean of all data points.

The neonatal Fc receptor (FcRn) is an MHC class I-like protein consisting of a heavy α chain encoded by the *FCGRT* gene, and a light β2-microglubulin (β2M) chain encoded by the *B2M* gene^15,16^, previously identified as a receptor for enteroviruses^17^. DPP4 is a cell surface exopeptidase important in glucose and insulin metabolism that also serves as the receptor for Middle East Respiratory Syndrome coronavirus^18,19^. Because both FcRn and DPP4 have been previously implicated as viral receptors, we sought to confirm the role of both factors in HAstV infection.

We disrupted *FCGRT, B2M,* and *DPP4* in Caco2 cells using two independent sgRNAs per gene and confirmed depletion of FcRn and DPP4 (**Supp Fig. 1B-E**). We found that HAstV1 and HAstV8 infection were significantly decreased when expression of either component of FcRn was disrupted (**Fig. 1B,C**). We stained Caco2 cells with anti-DPP4 antibody^18^ and observed DPP4 ubiquitously expressed in Caco2 cells using flow cytometry but found that HAstV-infected cells had higher levels of DPP4 compared to uninfected cells (**Fig. 1D**). As with FcRn, disruption of DPP4 also significantly reduced HAstV1 or HAstV8 infection (**Fig. 1E**). Disruption of *FCGRT* and *B2M* did not alter DPP4 expression, and correspondingly disruption of *DPP4* did not alter FcRn expression, indicating independent requirements for these factors for HAstV infection (**Supp Fig. 1B-D**). Pre-treatment of Caco2 cells with anti-DPP4 antibody effectively abrogated HAstV infection (**Fig. 1F and Supp Fig. 1F**), suggesting that DPP4 on the cell surface is crucial for HAstV entry. We confirmed that both factors are involved in viral entry by transfecting HAstV1 RNA into Caco2 cells disrupted for *DPP4* or *FCGRT* and demonstrating that viral protein expression proceeded similarly as in control Caco2 cells (**Supp Fig. 1G**).

### FcRn and DPP4 permit infection of non-permissive cells

In addition to our genome-wide CRISPR-Cas9 screen, we also performed a CRISPR activation (CRISPRa) screen, in which host factors are overexpressed instead of disrupted, as an alternative method for identifying important attachment and entry factors for HAstV. We leveraged a previously described “surfaceome” sgRNA library targeting 6,011 cell surface proteins ^20^ in Caco2 cells expressing catalytically “dead” Cas9 (dCas9) fused to the VP64 transcriptional activator to produce a pool of cells with induced expression of individual proteins. We infected this library with HAstV1 or HAstV8 at an MOI of 10 for 24 hours, sorted the top 3% of infected cells, and analyzed sgRNAs as in our genome-wide knockout screen. The top hit in the surfaceome screens for both HAstV1 and HAstV8 was *DPP4* (**Fig. 2A,B and Supp Fig. 2A**). To validate our screening results, we used CRISPRa to overexpress *DPP4* in Caco2 cells using two independent sgRNAs. We found that enhancement of DPP4 expression increased HAstV infection (**Fig. 2C and Supp Fig. 2B**). While *FCGRT* was one of the cellular surface factors assayed in the screen, it did not emerge as a hit. Assessment of the capacity of CRISPRa to induce FcRn overexpression using independent sgRNAs targeting *FCGRT* showed no effect, suggesting it may not have emerged from screening due to ineffective targeting by dCas9 (**Supp Fig. 2C**). While Caco2 cells support robust HAstV replication, HAstV replicates poorly in many routinely used immortalized cells including HEK293T ^21,22^. We assessed expression of DPP4 and FcRn in these cells and found that HEK293T express minimal FcRn and DPP4 (**Supp Fig. 2D,E**). We transiently transfected cDNA plasmids expressing human *FCGRT* or *DPP4* in HEK293T cells and found that overexpression of FCGRT or DPP4 was sufficient to permit HAstV1 infection (**Supp Fig. 2F,G**). We next generated HEK293T cell lines stably overexpressing FCGRT or DPP4 via lentiviral transduction. Stable overexpression of FCGRT promoted HAstV infection (**Fig. 2D and Supp Fig. 2D**). Similarly, HEK293T cells overexpressing DPP4 exhibited robust HAstV1 infection that could be blocked by anti-DPP4 antibody treatment (**Fig. 2E, F and Supp Fig. 2F**). Transient transfection of this HEK293T-DPP4 line to co-express FCGRT, but not B2M, with DPP4 exhibited increased HAstV1 infection (**Supp Fig. 2H**). Ectopic expression of DPP4 or FCGRT thus promotes HAstV infection in normally non-permissive cells, with enhanced activity when both factors are co-expressed, indicating that these receptors are sufficient for HAstV entry.

**Fig. 2.**
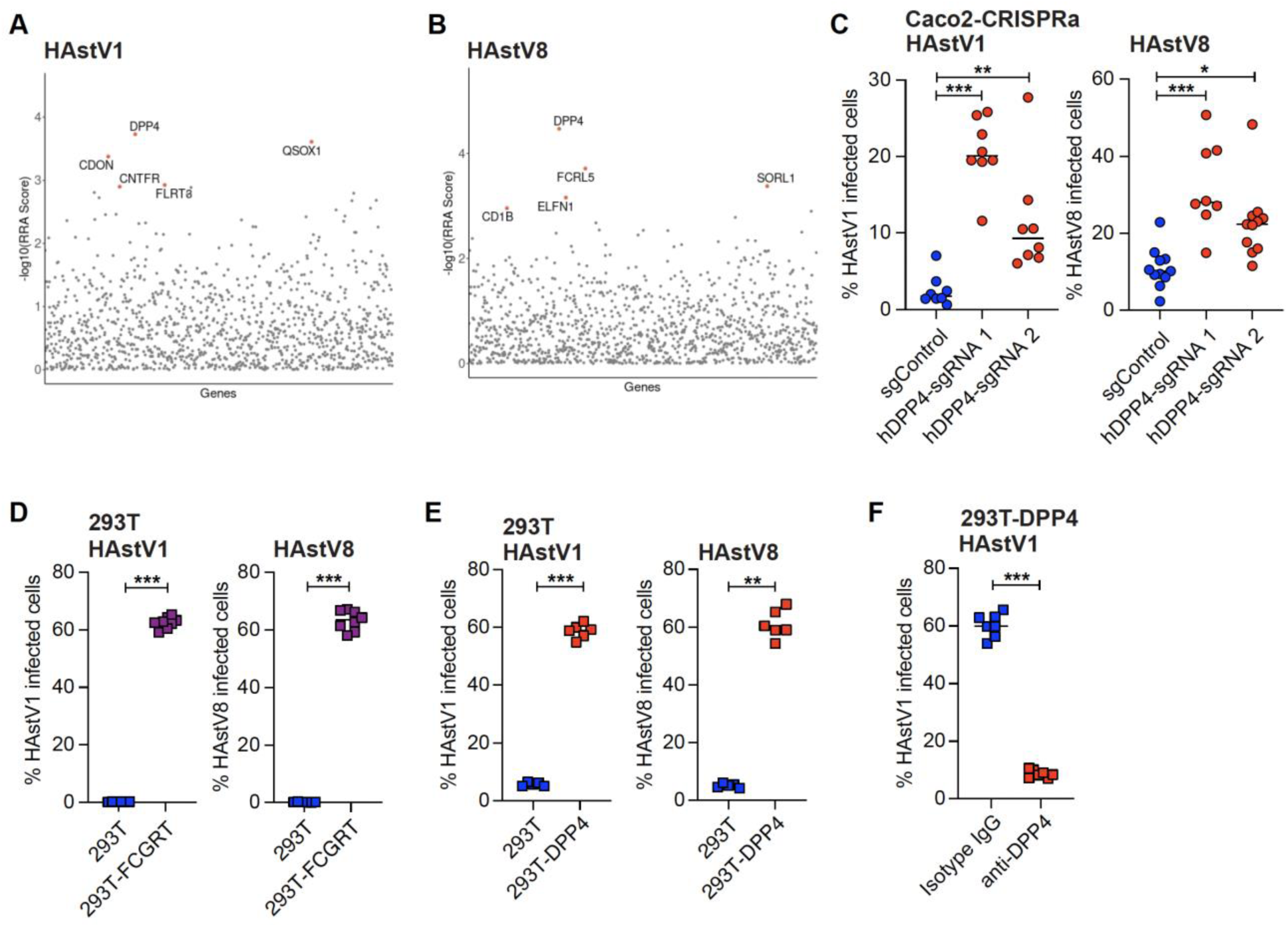
FcRn and DPP4 promote HAstV infection of non-permissive cells. (**A,B**) Enrichment scores of positively-selected sgRNAs in HAstV1-or HAstV8-positive cells sorted 24hpi from a surfaceome CRISPRa Caco2 cell library compared to unsorted cells, calculated by MAGeCK. (**C**) dCas9-Caco2 cells were targeted for *DPP4* (n=8) using two independent sgRNAs per gene or a control anti-GFP sgRNA (n=8), then assessed for infection using anti-HAstV capsid antibody 24hpi with HAstV1 or HAstV8. (**D, E**) HAstV1 levels in HEK293T (293T) (n=6) or HEK293T stably expressing FCGRT (293T-FCGRT) (n=8) or DPP4 (293T-DPP4) (n=6) at 24hpi. (**F**) HAstV1 levels in H293T-DPP4 cells treated with isotype control (n=7) or anti-DPP4 antibody (n=7) prior to HAstV infection at 24hpi. Results from three independent experiments were analyzed using Kruskal-Wallis test with Dunn’s post-test (C) or Mann-Whitney test (D to F). *P<0.05; **P<0.01; ***P<0.001; ****P<0.0001. Bars indicate mean of all data points.

### FcRn directly interacts with HAstV spike protein

To evaluate whether there are direct interactions between FcRn and the HAstV virion, we undertook surface plasmon resonance (SPR) experiments (**Fig. 3A**). Purified HAstV virion was flown over a biosensor with the recombinant ectodomains of FcRn immobilized using amine-coupling. The pan-HAstV mAb 8E7 that targets HAstV VP34 (**Fig. 3B**) was used as a positive control, while a chikungunya virus-specific mAb was used as a negative control. Our data showed that HAstV1 bound to both FcRn and mAb 8E7 to approximately 2000 response units (RU), and appeared to have a slow off-rate, which was expected and indicative of strong avidity. HAstV1’s interaction with immobilized FcRn had a similar binding curve to 8E7 with only a slightly faster off-rate.

**Fig. 3.**
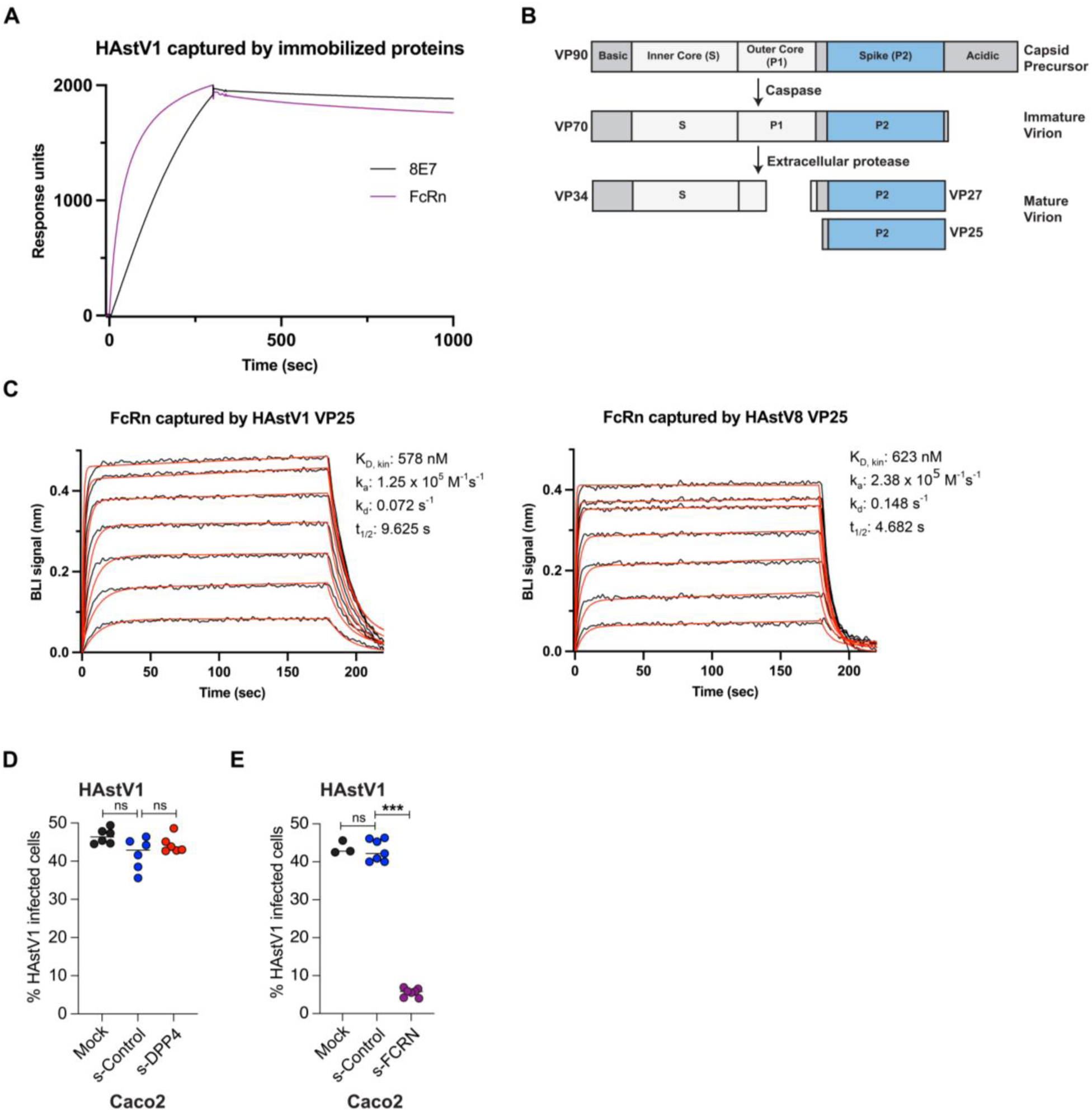
FcRn interacts with HAstV spike. (**A**) HAstV1 was tested against immobilized FcRn and mAb 8E7 via surface plasmon resonance. (**B**) Maturation process of HAstV VP90 structural protein. (**C**) VP25 protein from HAstV1 (left) and HAstV8 (right) was immobilized and tested against 2-fold dilutions of FcRn ranging from 8μM to 62.5nM via biolayer interferometry. (**D, E**) HAstV1 levels at 24hpi in Caco2 cells treated with PBS (Mock) (n=3-6), non-specific control protein (n=6-7), or soluble FcRN (s-FCRN) (n=7) or soluble DPP4 (s-DPP4) (n=6) prior to infection. Results from 2-3 independent experiments were analyzed using Kruskal-Wallis test with Dunn’s post-test (F and G). *P<0.05; **P<0.01; ***P<0.001; ****P<0.0001. ns=not significant. Bars indicate mean of all data points.

HAstV genomes are approximately 7kb with three open reading frames (ORFs); ORF2 encodes the structural capsid precursor protein VP90^23^ (**Fig. 3B**). This protein undergoes maturation via cleavage to produce the components of the mature virion: the capsid protein VP34 and the spike protein VP25/27. Previous antigenicity and sequence analyses suggest that the spike domain is likely to be involved in cell attachment and receptor binding^4–6^. We thus investigated if FcRn or DPP4 interact with the spike domain. We recombinantly expressed HAstV1 and HAstV8 spike proteins, VP25 (**Fig. 3B**), in *E. coli*, purified using both affinity- and size-exclusion chromatography (**Supp Fig. 3A,B**), and biotinylated the spike proteins. We also recombinantly expressed HAstV1 VP34 in Expi293 cells, purified using both affinity- and size-exclusion chromatography (**Supp Fig. 3C**). We generated the soluble extracellular domain of DPP4, confirmed that it was a dimer using MALS, and successfully tested the protein against a known binding partner, MERS-CoV spike^19^ (**Supp Fig. 3D,E**). We performed bio-layer interferometry (BLI) by immobilizing the spike proteins via streptavidin biosensors and dipped them into wells containing a range of concentrations of either FcRn or DPP4. FcRn bound to HAstV1 VP25 with an affinity of 578 nM and a half-life of 9.625 s, and FcRn bound to HAstV8 VP25 with a similar affinity of 623nM and a half-life of 4.682 s (**Fig. 3C**). DPP4 did not appear to interact with VP25 from either HAstV (**Supp Fig. 3F**).

To further understand the interaction of DDP4 with HAstV, we immobilized HAstV1 VP34 via anti-polyhistidine biosensors, and dipped into a range of concentrations of DPP4 (**Supp Fig. 3G**). Again, DPP4 did not appear to interact with recombinant VP34. The lack of observable interaction between DPP4 and the components of HAstV1 could be due to several circumstances. Prior cryogenic electron microscopy studies show that the mature virion has T =3 icosahedral symmetry containing a groove across the twofold axis and depressions at the fivefold and threefold symmetry axes, made by the interactions of the VP34 proteins^24^. It is possible that DPP4 targets a binding site specific to this arrangement of VP34 on the virion surface, and that recombinant VP34 cannot replicate this structure. Another possibility is that DPP4 requires a cofactor that is unaccounted for in these experiments. Finally, it is possible that expression of the soluble extracellular domain of DPP4 with a tag at the C-terminal domain affects its structure and function, as well as its ability to adequately interact with HAstV. As a Type II membrane protein, DPP4’s N-terminal domain is cytosolic and its C-terminal domain is extracellular. Using a tag at the C-terminal end of DPP4 to immobilize the protein via biosensors could obstruct its binding site and limit its interaction with HAstV. To determine whether soluble DPP4 or FcRn can neutralize HAstV infection in Caco2 cells, we incubated HAstV1 with soluble FcRn or DPP4 prior to infection and found that FcRn treatment, but not DPP4 treatment, was sufficient to prevent infection, suggesting that viral binding to cell membrane FcRn is a necessary step for HAstV infection (**Fig. 3D,E**).

### Pharmacological inhibition of FcRn or DPP4 prevents HAstV infection of *ex vivo* human enteroids

Numerous FcRn inhibitors, either modified IgG Fc fragments or anti-FcRn monoclonal antibodies with high affinity for FcRn that compete with IgG for binding to FcRn, are in clinical trials for their potential utility in treating myasthenia gravis and other autoimmune disorders^25^. We treated Caco2 cells with one of these antibodies, nipocalimab, and found that this treatment prevented HAstV1 infection (**Fig. 4A and Supp Fig. 4A**). Nipocalimab also blocked HAstV infection in FCGRT-expressing HEK293T cells (**Fig. 4B**).

**Fig. 4.**
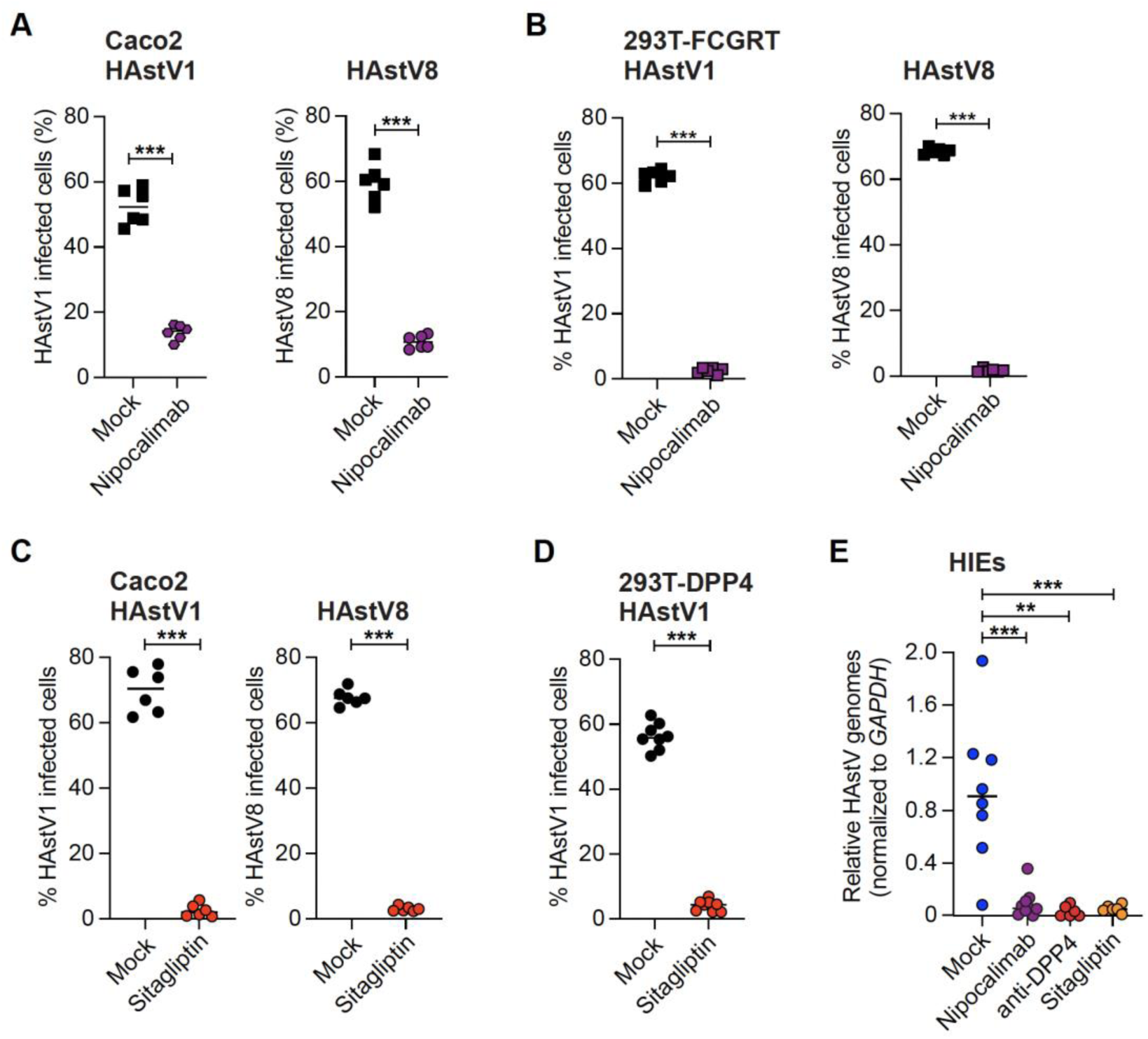
FcRn and DPP4 inhibitors are antiviral against HAstV. (**A,B**) HAstV1 and HAstV8 infection at 24hpi in Caco2 and 293T-FCGRT cells treated with PBS (n=6) or nipocalimab (n=6) prior to infection. (**C, D**) HAstV1 and HAstV8 infection at 24hpi in Caco2 or 293T-DPP4 cells treated with PBS (n=6-8) or sitagliptin (n=6-8) prior to infection. (**E**) HAstV1 levels at 24hpi in HIEs differentiated in monolayers on transwell inserts treated with PBS (n=7), nipocalimab (n=9), anti-DPP4 antibody (n=6) and sitagliptin (n=6) prior to infection. Results from three independent experiments were analyzed using Mann-Whitney test (A to D) or Kruskal-Wallis test with Dunn’s post-test (E). *P<0.05; **P<0.01; ***P<0.001; ****P<0.0001. Bars indicate mean of all data points.

Gliptins are a widely used class of oral medications to treat type 2 diabetes that specifically act to inhibit DPP4 enzymatic activity^26^, binding to the glucagon-like peptide interacting site. To evaluate if the enzymatic activity of DPP4 was necessary for HAstV entry, we administered sitagliptin, vildagliptin or teneligliptin to Caco2 cells prior to HAstV infection, using previously published dosing^18^. We found that while teneligliptin and vildagliptin exhibited minimal effects (**Supp Fig. 4B**), sitagliptin effectively blocked HAstV1 and HAstV8 infection of Caco2 cells (**Fig. 4C**). Similarly, HEK293T cells overexpressing DPP4 were sensitive to sitagliptin treatment (**Fig. 4D**). We confirmed inhibition of DPP4 enzymatic activity by all three gliptins (**Supp Fig. 4C**). These data suggest that the enzymatic activity of DPP4 is dispensable for HAstV infection, but also raise the possibility that sitagliptin may block DPP4-HAstV interactions via its specific mode of DPP4 binding.

HAstVs replicate robustly in human intestinal enteroids (HIEs)^27,28^. To evaluate the role of DPP4 and FcRn in this *ex vivo* system, we derived monolayer cultures from an ileal HIE line on transwells and infected them with HAstV1. Addition of nipocalimab, sitagliptin, or anti-DPP4 antibody prior to infection limited HAstV1 infection of these physiologically relevant cultures (**Fig. 4E**), underlining the potential for both these proviral factors as druggable targets in human-derived cells in addition to transformed cell lines.

Our work establishes that both FcRn and DPP4 are necessary and sufficient for HAstV entry and replication *in vitro* and can be readily targeted to prevent infection in *ex vivo* HIE cultures. The ability of anti-DPP4 antibody treatment to block HAstV infection while FcRn directly binds HAstV1 and 8 VP25 spike protein, indicates that DPP4 may serve as an initial factor critical for HAstV binding, while FcRn could be the receptor for HAstV entry. Indeed, our screening results paralleled a recent report identifying critical host factors for echovirus 6, in which *FCGRT*, *CD55*, and *B2M* emerged as the top three hits^17^. While CD55 serves as a critical receptor for echovirus attachment ^29^, FcRn is involved in echovirus uncoating and genome release^17,30^. HAstV may share similar entry mechanisms, with DPP4 replacing the activity of CD55. While cell surface DPP4 may bind HAstV capsid through other means, FcRn binds the spike protein, with both interactions ultimately required for replication. Our studies reveal sitagliptin, nipocalimab, and soluble FcRn as compelling antiviral candidates to prevent or limit HAstV infections, providing a path for therapeutic interventions against this diarrheal pathogen.

## Materials and Methods

### Cell culture

Caco2 and HEK293T cells were purchased from ATCC. Cells were cultured in Dulbecco’s Modified Eagle Medium (DMEM, Gibco #11995040) supplemented with 10% fetal bovine serum (FBS) and 1% penicillin and streptomycin (Gibco #15070063). Stable cell lines were generated via transduction as described previously^31^. For antibiotic selection, Caco2 medium was supplemented with 5μg/ml puromycin and 10μg/ml blasticidin, and HEK293T medium was supplemented with 1μg/ml puromycin or 50ug/ml neomycin sulfate when required.

### Gene cloning and protein purification

gBlocks (IDT technologies) containing the coding fragments corresponding to VP25 from HAstV1 (accession number AAC34717.1, residues S424 to R648) and HAstV8 (accession number AAF85964.1, residues S484 to R708) were cloned via Gibson assembly into a pET21 vector containing six histidine residues at the C-terminal end. They were then transformed into *Escherichia coli* strain BL21(DE3), and protein was expressed for 16 hours using autoinduction media at 18°C. Cells were lysed via ultrasonication in Buffer A (20mM Tris pH 8, 500mM NaCl, 20mM imidazole) with 2mM MgCl_2_, 0.25 mg/mL DNASE, and 0.25 mg/mL AEBSF, centrifuged, and the supernatant flowed over high-density nickel agarose beads (GoldBio H-320). After washing with Buffer A and eluting with 500mM imidazole, the proteins were further purified by flow over a HiLoad 16/600 Superdex 200 column in 20mM Tris pH 8, 150mM NaCl, and 0.1% sodium azide. The coding fragment corresponding to VP34 from HAstV1 (accession number above, residues V74 to G429) was inserted into a pCMV construct with an IL-2 signal sequence at the N-terminal end and a six-histidine tag at the C-terminal end. This was designed and synthesized by Twist Biosciences (San Francisco, CA). Constructs were transfected into Expi293 cells (Thermo Fisher) according to manufacturer’s instructions. Supernatants were collected four to five days post-transfection, centrifuged, and filtered. Supernatants were collected four to five days post-transfection, centrifuged, and filtered. Supernatants were flowed over high-density nickel agarose beads. After washing with 1X PBS and eluting with 500mM imidazole, the protein was dialyzed into 20mM HEPES, 150mM NaCl, and 0.1% sodium azide and further purified by flow over a Superdex 75 Increase 10/300 column. The coding fragment corresponding to the extracellular domain of human DPP4 (accession number NP_001926.2, residues S39-P766) was inserted into a pCMV construct with an IL-2 signal sequence at the N-terminal end and a twin-streptactin tag at the C-terminal end. This was designed and synthesized by Twist Biosciences and the construct was transfected into Expi293 cells according to manufacturer’s instructions. Supernatants were collected four to five days post-transfection, centrifuged, and filtered. Supernatants containing twin-streptactin-tagged DPP4 were flowed over Strep-Tactin XT resin (IBA Lifesciences), washed with 1X TBS, and eluted using the Strep-Tactin XT elution buffer (IBA Lifesciences). Protein was dialyzed into 20mM HEPES, 150mM NaCl, and 0.1% sodium azide and further purified over a Superdex 200 Increase 10/300 column. For MALS, recombinant DPP4 was loaded onto a size exclusion chromatography column in 20mM HEPES, 150mM HEPES pH 8, and 0.1% sodium azide in series with a MALS detector (DAWN) and an Optilab refractive index (RI) detector (Wyatt Technology). DPP4 was injected at ∼1mg/mL onto the WTC-015N5 (Wyatt Technology). Data was analyzed using Astra software (version 8.0.2.5).

### Virus infections and treatments

HAstV1 and HAstV8 were grown in Caco2 cells as described previously ^32^. For infections, 2.5×10^4^ Caco2 or HEK293T cells were seeded in 96-well plates 24h prior to infection. Cells were infected with HAstV at MOI 10 for 1h at 37°C, followed by 2 washes with PBS. Fresh media was added and the cells were incubated at 37°C until the end of the experiment. Cells were treated with 500μM sitagliptin (Sigma Aldrich; SML3205), 25μM vildagliptin (Sigma Aldrich; SML2302) and 20μM teneligliptin (Sigma Aldrich; SML3077) 12h prior to HAstV infection.

For DPP4 blocking experiments, cells were treated with 10μg/ml of goat IgG isotype control (Invitrogen, #02-6202) or human DPP4/CD26 antibody (R&D Systems; AF1180) 12h prior to HAstV infection. The treatments were maintained throughout the experiment. For FcRn blocking experiments, cells were treated with 10μg/ml nipocalimab (Medchem Express; HY-P99037) 12h prior to HAstV infection and maintained in media containing nipocalimab throughout the experiment.

### Generation of genetically modified cell lines

Individual sgRNAs for targeting the candidate genes were designed using the CRISPick program of the Broad Institute. These sgRNAs were cloned into linearized lentiGuide-Puro (Addgene # 52963) for gene deletion, and pXPR_502 (Addgene #96923) for CRISPRa. Lentivirus was generated by transfecting lentiviral vectors with packaging vector (psPax2; Addgene #12260) and pseudotyping vector (pCMV-VSV-G; Addgene #8454) into HEK293T cells using TransIT-LT1 (Mirus). Then, 48 h later, supernatants were filtered through a 0.45-μm pore filter and stored in - 80C until use. Caco2-Cas9 and Caco2-dCas9 cells were generated by transducing with pLX311-Cas9 (Addgene #118018) and pLenti-dCas9-VP64-Blast (Addgene #61425), respectively. After 10 days of blasticidin selection, cells were transduced with individual lentiviruses targeting the candidate genes by spin transduction. Cells were grown in blasticidin and puromycin for 7 days to select for transduced cells. Stable DPP4 expressing HEK293T cells were generated by transducing HEK293T cells with hDPP4-puro (Genscript) lentivirus followed by selection with puromycin for 7 days. Overexpression of DPP4 in stable cells was confirmed by flow cytometry. Stable FcRn expressing HEK-293 cells were generated by transfecting hFCGRT-neo plasmid (Genscript) in HEK293 cells using TransIT-LT1 (Mirus). After 48 h, cells were grown in neomycin sulfate for 10 days to select FcRn expressing stable cells. Overexpression of FcRn was confirmed in the stable cells by western immunoblotting analysis.

### CRISPR knockout screening

Human genome-wide CRISPR knockout pooled library (“Brunello”) was a gift from David Root and John Doench (Addgene #73178)^10^. Caco2-Cas9 cells were transduced with this library at approximately 0.3 MOI to obtain 40 million transduced cells, which is sufficient for integration of each sgRNA into approximately 500 cells. At 2d post-transduction puromycin was added and cells were selected for at least 10 days. For screening, 1×10^7^ cells per sample were plated in a 15cm^2^ culture dish. After 24 h, the cells were infected with HAstV1 or HAstV8 at MOI 10 for 24h. The cells were washed with PBS twice to remove unbound virus and detached using trypsin-EDTA. The cells were fixed and permeabilized using BD Cytofix/Cytoperm (BD Biosciences) buffer followed by washing with BD Perm/Wash buffer (BD Biosciences). Then, the cells were incubated with 10ug/ml of astrovirus type 2 monoclonal antibody (8E7) (Invitrogen, USA) overnight. After washing 2 times with BD Cytofix/Cytoperm (BD Biosciences) buffer, cells were stained with PE-conjugated anti-mouse IgG antibody (Jackson ImmunoResearch) for 1 h at room temperature followed by washing 2 times with BD Cytofix/Cytoperm buffer. The cells were subjected to sorting using FACS AriaIII (BD Biosciences) to collect PE-negative (uninfected) cells as well as PE-positive (infected) cells. Genomic DNA (gDNA) was extracted from the isolated cells and unsorted cells (input) with QIAamp DNA Maxi kit (Qiagen; #51104).

### CRISPR activation screening

A list of 1100 surface proteins was obtained based on datasets for plasma membrane proteins ^33^. Four sgRNAs targeting each gene were picked from Calabrese genome-wide CRISPRa library, and 250 nontargeting control sgRNAs were included. The sgRNAs were cloned into pXPR_502 (Addgene, #96923) with assistance from the Genome Engineering and iPSC Center (GEiC) at Washington University in Saint Louis. Forty million Caco2-dCas9 cells were transduced with the CRISPRa library at approximately 0.3 MOI to get 10 million transduced cells, which is sufficient for the integration of each sgRNA into approximately 1000 cells. At 2 d post-transduction puromycin was added and cells were selected for at least 10 days. For screening, cells were plated, infected, and stained as for CRISPR knockout screening, with screening performed for either HAstV1 or HAstV8. Cells were sorted using FACS to collect the top 3% (fluorescence intensity) of infected cells. As a control, the same number of cells were stained with PE anti-CD45 antibody (BioLegend; #368522) and the top 3% of PE-positive cells were sorted. Genomic DNA (gDNA) was extracted from the isolated cells and unsorted cells (input) with QIAamp DNA Maxi kit (Qiagen; #51104).

### Sequencing and analysis

For Illumina sequencing, gDNA was used for PCR to amplify the integrated sgRNA sequences. PCR was performed in 96-well plates and each well containing up to 10 μg of gDNA in a total of 100 μL reaction mixture consisting of Titanium Taq DNA polymerase buffer and enzyme (Takara, #639209), deoxynucleoside triphosphate, dimethylsulfoxide (5%), P5 stagger primer mix (0.5 μM), and uniquely barcoded P7 primer (0.5 μM). Samples were amplified with following PCR cycles: an initial 5 min at 95°C; followed by 35 cycles of 95°C for 30 s, 59°C for 30 s, and 72°C for 20 s; followed by a final 10 min at 72°C. PCR products were pooled and purified with AMPure XP beads according to the manufacturer’s protocol (Beckman Coulter, #MSPP-A63880). Samples were sequenced on a MiSeq sequencer (Illumina). After demultiplexing according to the barcode sequences, reads were mapped to a reference file of sgRNAs in CRISPR library using MAGeCK Version 0.5.9^14^. We calculated a Robust Ranking Algorithm (RRA) score of sgRNAs in each sample relative to control (infected cells for CRISPR KO screen, and unsorted cells in CRISPRa screen) to identify CRISPR-altered genes overrepresented in the populations of interest.

### Western blotting

For western blotting, cells were lysed directly in Laemmli buffer, denatured by boiling, and subjected to SDS-PAGE using Bio-Rad Any kD™ Mini-PROTEAN® TGX gels. Protein was transferred to PVDF membrane and stained using anti-FCGRT polyclonal antibody (Invitrogen PA5-97738) and subsequently DyLight 800-conjugated anti-rabbit secondary antibody (Invitrogen SA5-10036). Loading control was performed using rhodamine-conjugated anti-actin fAB fragment (Bio-Rad 12004163). Blots were imaged using a Bio-Rad Chemidoc fluorescent imaging device.

### Neutralization assay

To assess the neutralization of HAstV infection by soluble DPP4 and soluble FcRn, 4×10^4^ FFU/ml of HAstV1 were incubated with 5μg/ml recombinant human DPP4/CD26 protein (Acro Biosystems; DP4-H5266) or 5μg/ml recombinant FCGRT/B2M complex (BPS Bioscience, 71283-1) in growth media at 37°C. Cells grown in 96-well plates were treated with this complex at 37°C for 1h, followed by PBS wash. The cells were grown for 24h to determine HAstV infection neutralization by flow cytometry assay.

### Flow cytometry

To assess HAstV infection, cells were fixed and permeabilized with BD Cytofix/Cytoperm for 15 min at RT, followed by two washes with BD Perm/Wash buffer. Cells were incubated with 1:1000 dilution of astrovirus type 2 monoclonal antibody (8E7) (Invitrogen, USA) for 1h at RT or overnight at 4°C, followed by two washes with BD Perm/Wash buffer. Next, the cells were incubated with 1:1000 dilution of PE-conjugated anti-mouse IgG antibody for 1h at RT in dark. Cells were washed twice and resuspended in BD Perm/Wash buffer, and analyzed by flow cytometry using BD FACS Canto II (BD Biosciences). Staining for DPP4 was performed with anti-DPP4 antibody (R&D Systems; AF1180) at 1:500 dilution and Alexa 647-conjugated anti-goat IgG secondary antibody (Jackson Immunoresearch; 705-605-003) at 1:1000 with the same incubation times and buffers. Data were processed using FACSDiva software (BD Biosciences) and FlowJo (BD Biosciences).

### Surface plasmon resonance (SPR)

SPR binding experiments were performed using the Biacore T200 system (GE Healthcare). Experiments were performed at 25μl min^−1^ and 25°C using 1X HBS-P buffer (10mM HEPES pH 7.6, 150mM NaCl, 0.05% (v/v) Tween-20). FcRn (R&D Systems 8639-FC) and 8E7 (Bio-Rad MCA2716) were immobilized onto a CM5 sensor chip to approximately 6000 response units (RU) using standard amine coupling. Purified HAstV1 was buffer exchanged into running buffer and concentrated before injection over the immobilized proteins. All results were reference-subtracted using the reference flow cell (immobilized CHKV.124.A4.F12 mAb, unpublished). All results were analyzed using BIAevaluation v.3.1 (GE Healthcare).

### Bio-layer interferometry (BLI)

BLI binding experiments were performed using the GatorPlus BLI system (GatorBio). Experiments were performed with 1X HBS-P buffer (10mM HEPES pH 7.6, 150mM NaCl, 0.05% (v/v) Tween-20) supplemented with 1% (w/v) BSA. Recombinant VP25s from both HAstV1 and HAstV8 were biotinylated using EZ-link NHS-PEG4-Biotin (Thermo Fisher) according to manufacturer’s instructions. The biotinylated proteins were immobilized on streptavidin biosensors (GatorBio 160002) and tested against two-fold serial dilutions of either FcRn (8μM to 62.5nM) or DPP4 (1μM to 15.625nM). Recombinant MERS-CoV spike and recombinant VP34 were immobilized on anti-his biosensors (GatorBio 160009) and tested against two-fold serial dilutions of DPP4 from 1μM to 62.5nM. All kinetic profiles were fitted using a global 1:1 binding algorithm with a drifting baseline using BIAevaluation v.3.1 (GE Healthcare).

### Human intestinal enteroid culture

Human intestinal tissue biopsies were obtained with approval from the Washington University in St. Louis institutional review board (no. 201804040). The HIEs were generated as described previously^34^. Briefly, tissue sample was digested in collagenase I (Invitrogen, Waltham, MA) for 10 min at 37°C followed by mechanical dissociation. The resulting crypts were filtered through a 70 μm cell strainer and washed in Dulbecco’s modified Eagle medium (Gibco) supplemented with 10% fetal bovine serum before resuspending in growth factor-reduced Matrigel (Becton Dickinson, Franklin Lakes, NJ), and seeded into culture plates. Enteroids were grown in 50% L-WRN conditioned medium supplemented with 10 μM Y-27632 and SB 202190 (R&D Systems). For transwell monolayer cultures, small intestinal enteroids were dissociated into single cells by TrypLE digestion prior to seeding into transwells with 0.4 μm pore size (Corning) precoated with 10% Matrigel (Corning) for 15 min^35,36^. Cells were cultured in 50% L-WRN conditioned media with 10 μM Rock inhibitor Y-27632 (R&D Systems). After 4 days of culture, the media was switched to differentiation media (5% L-WRN conditioned). Monolayers were then grown in differentiation media for up to 4 days before initiating experiments. Media was changed every 1–2 days. Sitagliptin, human DPP4 antibody and nipocalimab were administered apically in the differentiation media for 12h prior to infection. HAstV infections were performed apically in serum free media at 37°C for 1h. Monolayers were maintained in differentiation media containing treatments.

### RNA extraction and quantitative reverse transcription PCR

Total RNA from tissues or cells was isolated using Tri Reagent (Invitrogen) and a Direct-zol-96 RNA kit (Zymo Research) according to the manufacturer’s protocol. ImPromII reverse transcriptase was used for cDNA synthesis (Promega). SYBR green qPCR assay was performed for HAstV1 using primers specific for HAstVs [Mon269 (5′-CAACTCAGGAAACAGGGTGT-3′) and Mon270 (5′-TCAGATGCATTGTCATTGGT-3′)]^28,37^. Quantitative polymerase chain reaction (PCR) for housekeeping gene GAPDH was performed with forward primer 5′-GACCTGCCGTCTAGAAAAACC-3′, reverse primer 5′-GCTGTAGCCAAATTCGTTGTC-3′, (Integrated DNA Technologies). SYBR green qPCR assays were performed with Power SYBR Green PCR master mix (ThermoFisher).

### DPP4 activity assay

Caco2 cells were plated in 6-well plates at 2 million cells per well and treated for 24h with 500μM sitagliptin, 25μM vildagliptin, or 20μM, then lysed and processed using DPP4 Activity Assay Kit (Sigma MAK088). After treatment with DPP4 substrate H-Gly-Pro-AMC, samples were incubated for 10 min at 37°C and AMC fluorescence was measured with 360 nm excitation and 460 nm emission.

## Acknowledgments

We thank all members of the Baldridge laboratory for helpful discussions. The human Brunello CRISPR knockout pooled library was a gift from David Root and John Doench (Addgene #73178). We additionally thank the Precision Animal Models and Organoids core for providing human intestinal organoid lines.

## Author contributions

H.I., J.M.M, P.D.H, D.B., L.W., and L.F performed the experiments. H.I., J.M.M, P.D.H, D.B., L.W. D.H.F., and M.T.B. analyzed the data. H.I., J.M.M, P.D.H., D.H.F., and M.T.B designed the project. A.A. and S.D. provided critical reagents. H.I., J.M.M. and M.T.B wrote the manuscript. All authors read and edited the manuscript.

## Data and materials availability

All data needed to evaluate the conclusions in the paper are present in the paper and/or the Supplementary Materials. CRISPR screening data are available at European Nucleotide Archive, accession PRJEB71796. CRISPR libraries and cell lines can be provided by Dr. Baldridge at Washington University School of Medicine pending scientific review and a completed material transfer agreement. Requests for these materials should be submitted to.

## Supplementary Materials

**Fig. S1.**
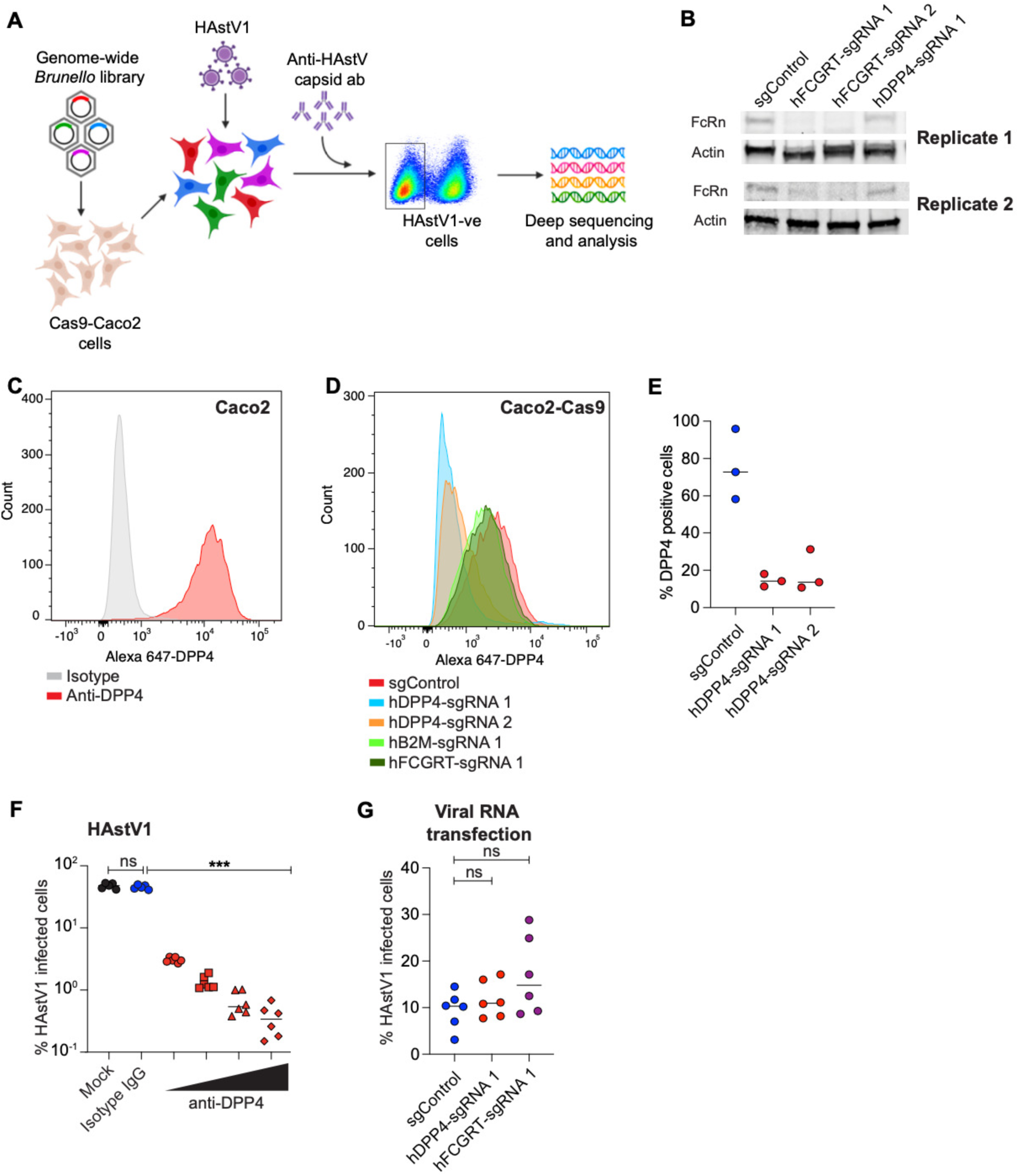
Disruption of FcRn or DPP4 in Caco2 cells limits HAstV infection. (**A**) Overview of the pooled CRISPR screen. The genome-scale Brunello library was introduced into Cas9 expressing Caco2 cells, followed by selection of transduced cells. After 10 days, pooled Caco2 cells were infected with HAstV1 for 24h, followed by staining using anti-HAstV capsid antibody. Uninfected cells were sorted to determine the sgRNA counts by next-generation sequencing. (**B**) FcRn protein levels in Cas9-Caco2 cells disrupted for FCGRT or DPP4 using independent sgRNAs per gene or targeted with a control anti-GFP sgRNA. Two independent replicates are shown. (**C**) Representative histogram for surface expression of DPP4 in naïve Caco2 cells stained with anti-DPP4 antibody or isotype control antibody. (**D**) Representative histogram showing DPP4 expression in fixed, permeabilized Caco2 cells disrupted for FCGRT or DPP4 or B2M using gene specific sgRNAs or targeted with a control anti-GFP sgRNA. (**E**) Percentage of DPP4 expressing Caco2 cells disrupted for DPP4 using sgRNAs or targeted with a control anti-GFP sgRNA. (n=3) (**F**) Percentage of anti-HAstV capsid antibody stained Caco2 cells treated with PBS (n=5) or isotype antibody (n=5) or various concentrations of anti-DPP4 antibody (n=6) prior to HAstV1 infection. (**G**) Percentage of anti-HAstV capsid antibody-stained control (n=6), FCGRT (n=6) or DPP4 (n=6) knockout Caco2 cells transfected with HAstV1 RNA at 48h post-transfection. Results were analyzed using Kruskal-Wallis test with Dunn’s post-test (F and G) from two to three independent experiments. *P<0.05; **P<0.01; ***P<0.001; ****P<0.0001. ns=not significant. Bars indicate mean of all data points.

**Fig. S2.**
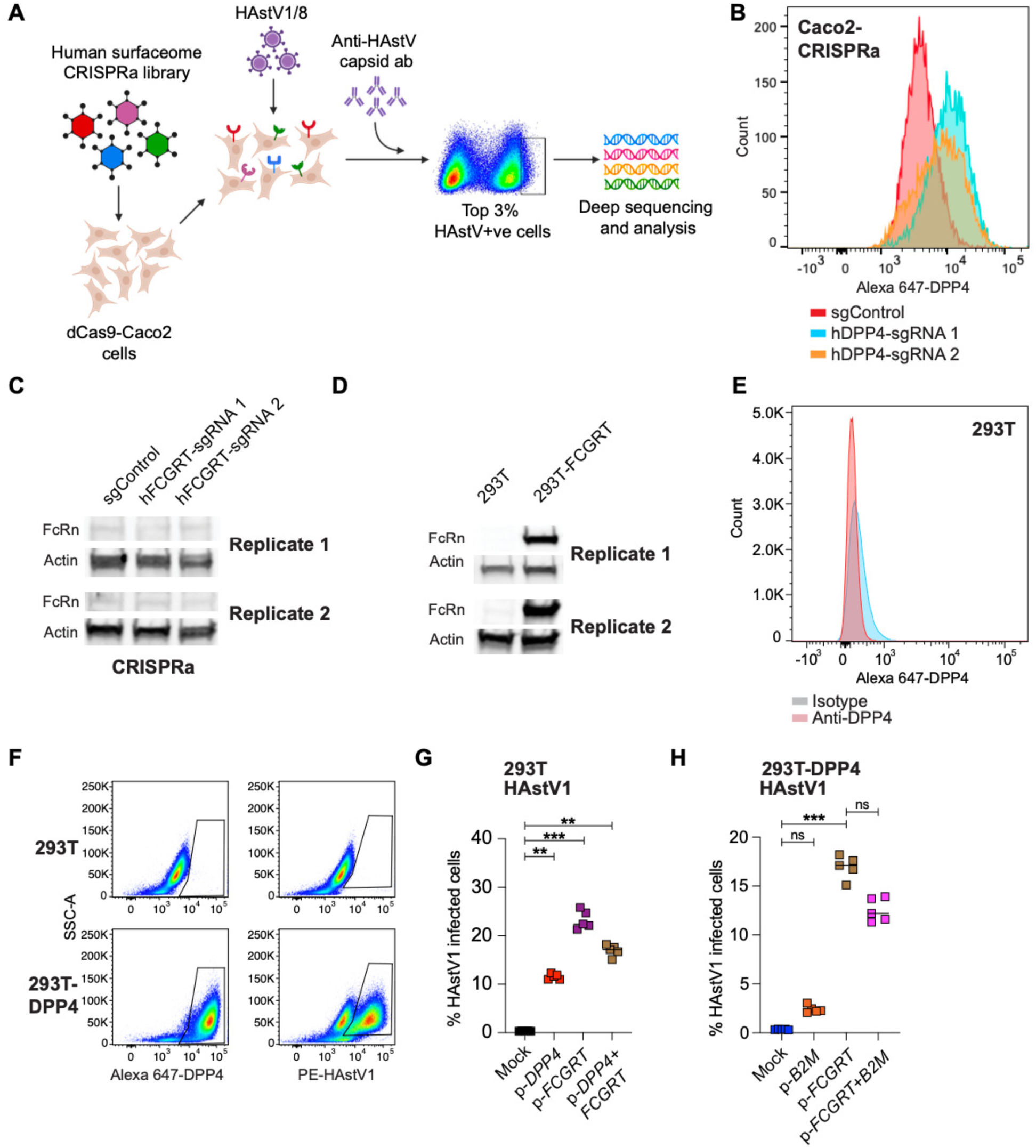
Enhanced expression of DPP4 or FCGRT promotes HAstV infection. (**A**) Schematic of the CRISPR activation surfaceome screen. The human surfaceome library was introduced into dCas9 expressing Caco2 cells, followed by selection of transduced cells. After 10 days, pooled Caco2 cells were infected with HAstV1 or HAstV8 for 24h, followed by staining using anti-HAstV capsid antibody. We sorted top 3% of the HAstV capsid positive cells to determine the sgRNA counts by next-generation sequencing. (**B**) Representative histogram showing expression of DPP4 in dCas9-Caco2 cells transduced with sgRNAs for DPP4 overexpression. (**C**) FcRn protein levels in dCas9-Caco2 transduced with sgRNA for overexpressing FcRn. Two independent replicates are shown. (**D**) FcRn protein levels in 293T and 293T-FCGRT cells. Two independent replicates are shown. (**E**) Representative histogram showing surface expression of DPP4 in 293T cells stained with anti-DPP4 antibody or isotype control antibody. (**F**) Abundance of DPP4 and HAstV capsid positive 293T and 293T-DPP4 cells infected with HAstV1. (**G, H**) HEK293T or HEK293T-DPP4 cells were transfected with *FCGRT* (n=5)*, DPP4* (n=5) and/or *B2M* (n=5) plasmids then 48h later were infected and stained with anti-HAstV capsid antibody at 24 hpi. Results were analyzed using Kruskal-Wallis test with Dunn’s post-test (G and H) from three independent experiments. *P<0.05; **P<0.01; ***P<0.001; ****P<0.0001. ns=not significant. Bars indicate mean of all data points.

**Fig. S3.**
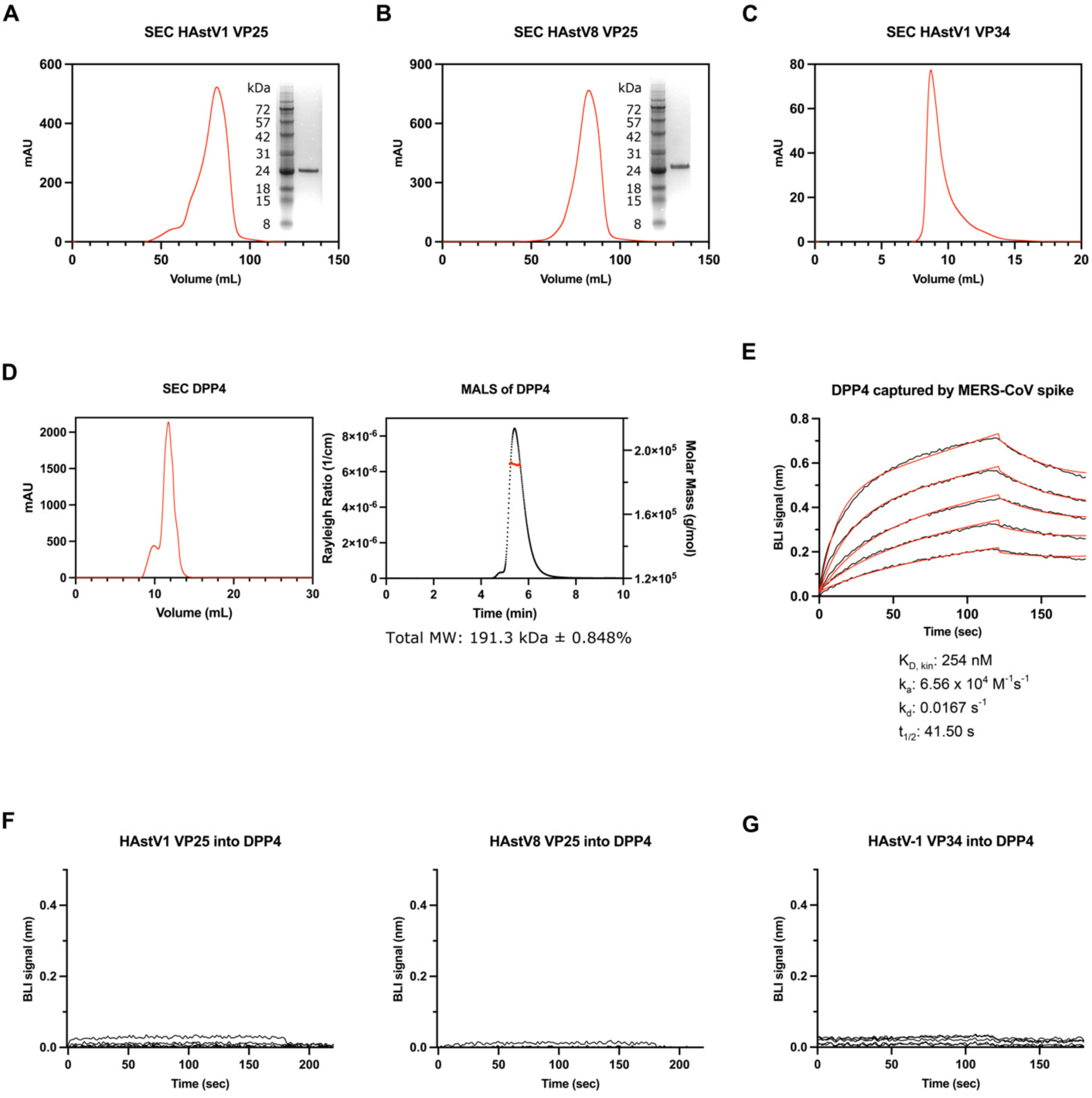
Purified DPP4 does not directly bind VP25 or VP34. (**A-C**) Gel filtration and SDS-PAGE profiles of HAstV1 VP25 (**A**) HAstV8 VP25 (**B**) and HAstV1 VP34 (**C**). HAstV1 and HAstV8 VP25 were eluted from a HiLoad 16/600 Superdex 200 column and HAstV1 VP34 was eluted from a Superdex 75 Increase 10/300 column. (**D**) Soluble hDPP4 was expressed via transfection in Expi293 cells and eluted via gel filtration on Superdex 200 Increase 10/300 column (left) and further analyzed using MALS (right). The MALS curve (black) is plotted with the derived molecular weight (red) of 191,3000 Dalton ± 0.848%. (**E-G**) Binding experiments testing full-length spike protein from MERS-CoV, (**E**) VP25 from HAstV1 and HAstV8 (**F**, left and right, respectively), and VP34 from HAstV1 (**G**) against DPP4. In all cases, each protein was immobilized via biosensor and tested against 2-fold dilutions of DPP4 from either 1μM to 62.5nM (**E, G**) or 1μM to 15.625nM (**F**).

**Fig. S4.**
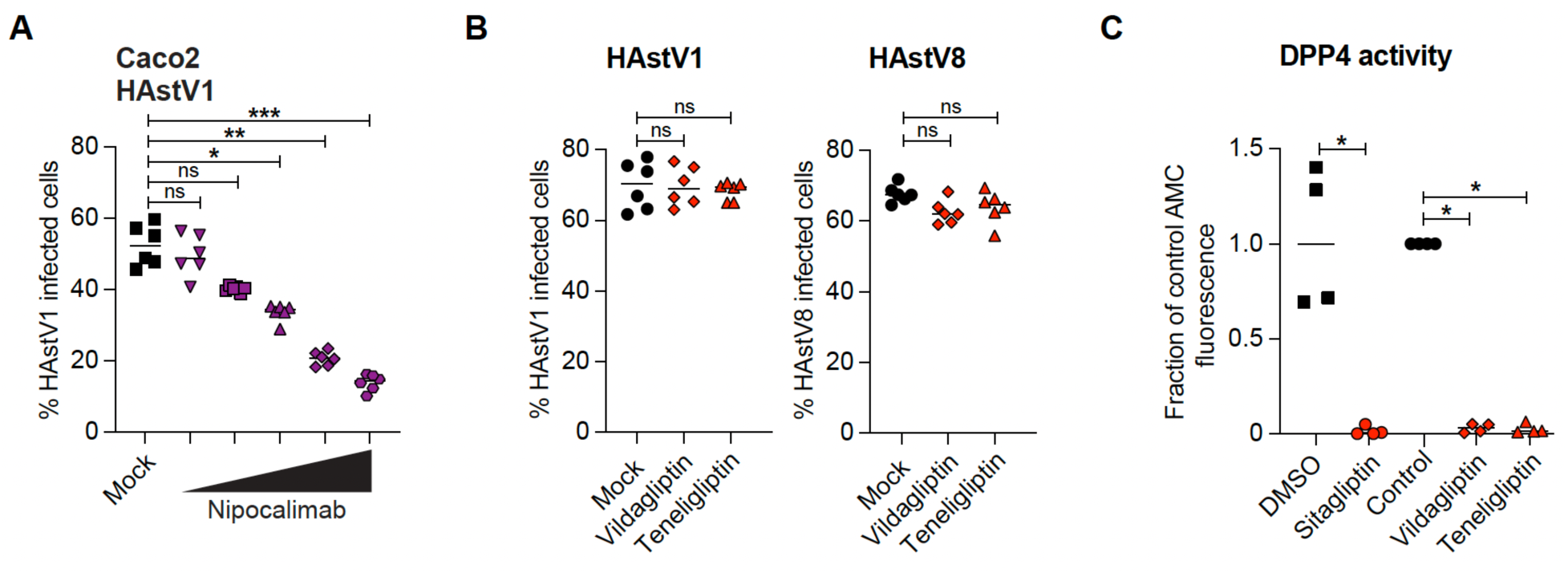
Despite similar inhibition of DPP4 activity, vildagliptin and teneligliptin do not prevent HAstV infection. (**A**) Percentage of anti-HAstV capsid antibody-positive Caco2 cells at 24hpi treated with PBS (n=6) or various concentrations of nipocalimab (n=6) prior to HAstV1 infection. (**B**) Percentage of anti-HAstV capsid antibody-positive Caco2 cells treated with PBS (n=6) or vildagliptin (n=6) or teneligliptin (n=6) prior to HAstV1 or HAstV8 infection. (**C**) Fluorescent 7-Amino-4-Methyl Coumarin (AMC) released by DPP4 enzymatic activity in Caco2 cells treated with PBS (Control; n=4) or DMSO or DPP4 inhibitors (n=4). Results were analyzed using Kruskal-Wallis test with Dunn’s post-test (**A**) from three independent experiments. *P<0.05; **P<0.01; ***P<0.001; ****P<0.0001. ns=not significant. Bars indicate mean of all data points.

### Flow cytometry gating strategy

Cells were gated on FSC/SSC to select the entire population of cells, then gated on FSC-A/FSC-H to select for single cells. Flow plots shown reflect the entire cell population without subgating. To assess for HAstV infection, gating for positive vs. negative cells was done using uninfected vs. infected control cells and subsequently applied uniformly to experimental cells to avoid bias.

