## Supplementary Data for "The neonatal Fc receptor and DPP4 are human astrovirus receptors"

**This file includes:**

Figs. S1 to S4

Flow cytometry gating strategy

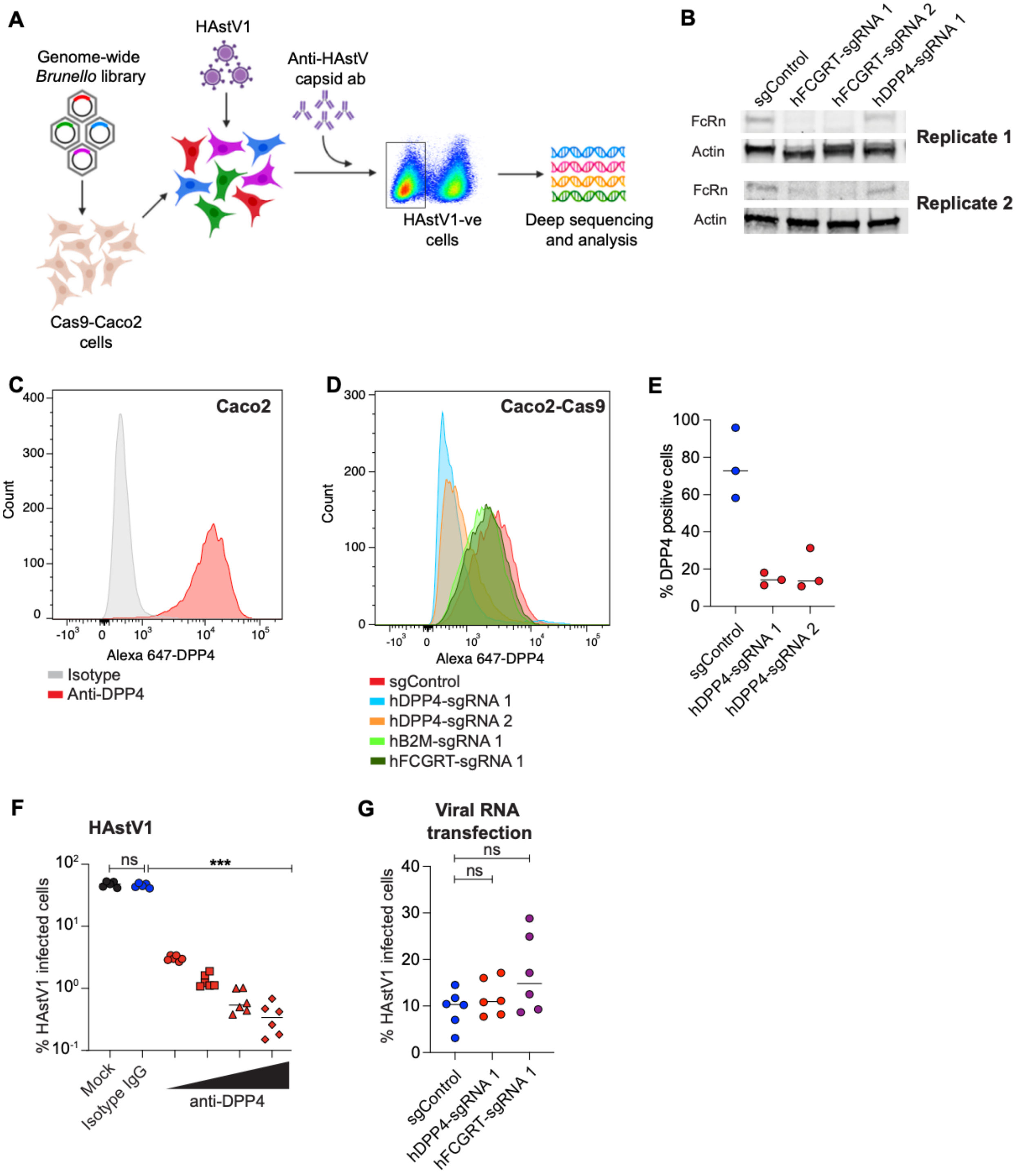


**Fig. S1. Disruption of FcRn or DPP4 in Caco2 cells limits HAstV infection.** (**A**) Overview of the pooled CRISPR screen. The genome-scale Brunello library was introduced into Cas9 expressing Caco2 cells, followed by selection of transduced cells. After 10 days, pooled Caco2 cells were infected with HAstV1 for 24h, followed by staining using anti-HAstV capsid antibody. Uninfected cells were sorted to determine the sgRNA counts by next-generation sequencing.

(**B**) FcRn protein levels in Cas9-Caco2 cells disrupted for FCGRT or DPP4 using independent sgRNAs per gene or targeted with a control anti-GFP sgRNA. Two independent replicates are shown. (**C**) Representative histogram for surface expression of DPP4 in naïve Caco2 cells stained with anti-DPP4 antibody or isotype control antibody. (**D**) Representative histogram showing DPP4 expression in fixed, permeabilized Caco2 cells disrupted for FCGRT or DPP4 or B2M using gene specific sgRNAs or targeted with a control anti-GFP sgRNA. (**E**) Percentage of DPP4 expressing Caco2 cells disrupted for DPP4 using sgRNAs or targeted with a control anti-GFP sgRNA. (n=3) (**F**) Percentage of anti-HAstV capsid antibody stained Caco2 cells treated with PBS (n=5) or isotype antibody (n=5) or various concentrations of anti-DPP4 antibody (n=6) prior to HAstV1 infection. (**G**) Percentage of anti-HAstV capsid antibody-stained control (n=6), FCGRT (n=6) or DPP4 (n=6) knockout Caco2 cells transfected with HAstV1 RNA at 48h post-transfection. Results were analyzed using Kruskal-Wallis test with Dunn's post-test (F and G) from two to three independent experiments. *P<0.05; **P<0.01; ***P<0.001; ****P<0.0001. ns=not significant. Bars indicate mean of all data points.

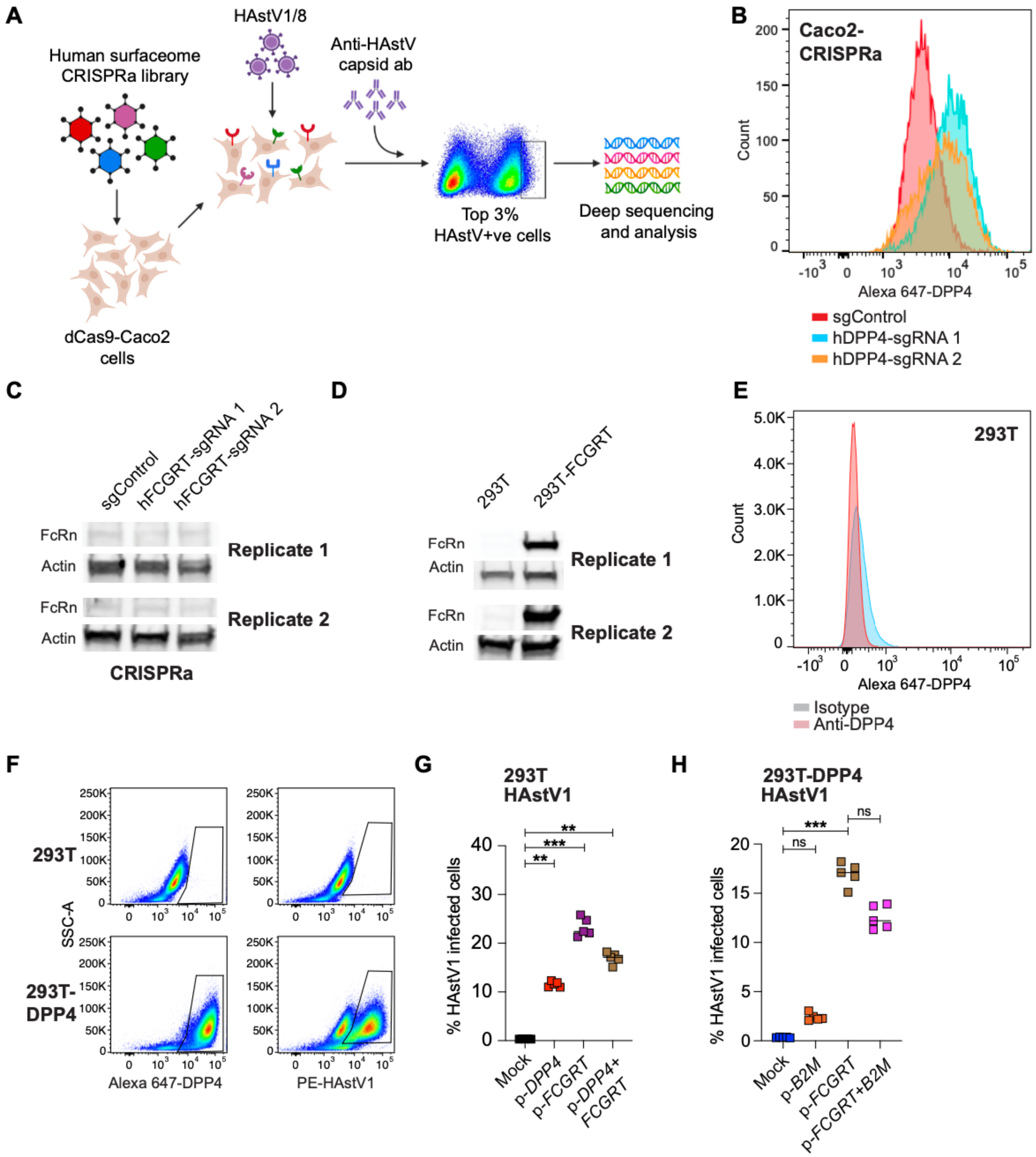


**Fig. S2. Enhanced expression of DPP4 or FCGRT promotes HAstV infection.** (**A**) Schematic of the CRISPR activation surfaceome screen. The human surfaceome library was introduced into dCas9 expressing Caco2 cells, followed by selection of transduced cells. After 10 days, pooled Caco2 cells were infected with HAstV1 or HAstV8 for 24h, followed by staining using anti-HAstV capsid antibody. We sorted top 3% of the HAstV capsid positive cells to determine the sgRNA counts by next-generation sequencing. (**B**) Representative histogram showing expression of DPP4 in dCas9-Caco2 cells transduced with sgRNAs for DPP4 overexpression. (**C**) FcRn protein levels in dCas9-Caco2 transduced with sgRNA for overexpressing FcRn. Two independent replicates are shown. (**D**) FcRn protein levels in 293T and 293T-FCGRT cells. Two independent replicates are shown. (**E**) Representative histogram showing surface expression of DPP4 in 293T cells stained with anti-DPP4 antibody or isotype control antibody. (**F**) Abundance of DPP4 and HAstV capsid positive 293T and 293T-DPP4 cells infected with HAstV1. (**G, H**) HEK293T or HEK293T-DPP4 cells were transfected with *FCGRT* (n=5), *DPP4* (n=5) and/or *B2M* (n=5) plasmids then 48h later were infected and stained with anti-HAstV capsid antibody at 24 hpi. Results were analyzed using Kruskal-Wallis test with Dunn's post-test (G and H) from three independent experiments. *P<0.05; **P<0.01; ***P<0.001; ****P<0.0001. ns=not significant. Bars indicate mean of all data points.

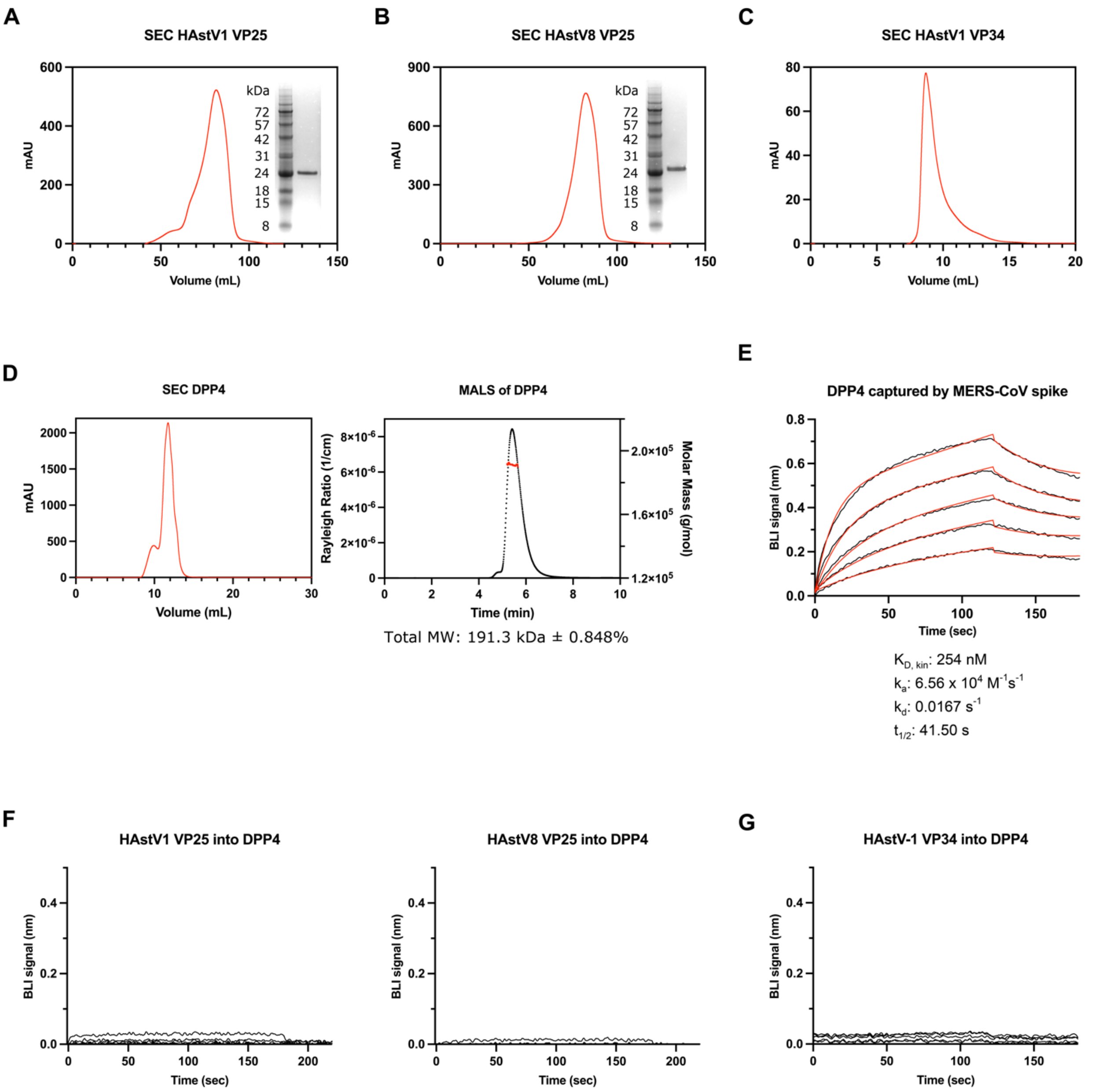


**Fig. S3. Purified DPP4 does not directly bind VP25 or VP34.** (**A-C**) Gel filtration and SDS-PAGE profiles of HAstV1 VP25 (**A**) HAstV8 VP25 (**B**) and HAstV1 VP34 (**C**). HAstV1 and HAstV8 VP25 were eluted from a HiLoad 16/600 Superdex 200 column and HAstV1 VP34 was eluted from a Superdex 75 Increase 10/300 column. (**D**) Soluble hDPP4 was expressed via transfection in Expi293 cells and eluted via gel filtration on Superdex 200 Increase 10/300 column (left) and further analyzed using MALS (right). The MALS curve (black) is plotted with the derived molecular weight (red) of 191,3000 Dalton ± 0.848%. (**E-G**) Binding experiments testing full-length spike protein from MERS-CoV, (**E**) VP25 from HAstV1 and HAstV8 (**F**, left and right, respectively), and VP34 from HAstV1 (**G**) against DPP4. In all cases, each protein was immobilized via biosensor and tested against 2-fold dilutions of DPP4 from either 1μM to 62.5nM (**E, G**) or 1μM to 15.625nM (**F**).

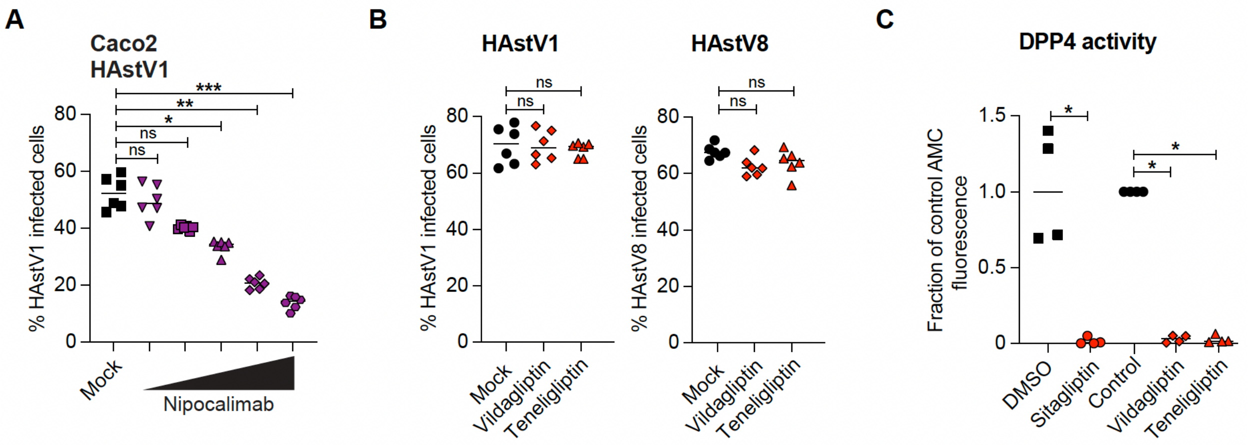


**Fig. S4. Despite similar inhibition of DPP4 activity, vildagliptin and teneligliptin do not prevent HAstV infection.** (**A**) Percentage of anti-HAstV capsid antibody-positive Caco2 cells at 24hpi treated with PBS (n=6) or various concentrations of nipocalimab (n=6) prior to HAstV1 infection. (**B**) Percentage of anti-HAstV capsid antibody-positive Caco2 cells treated with PBS (n=6) or vildagliptin (n=6) or teneligliptin (n=6) prior to HAstV1 or HAstV8 infection. (**C**) Fluorescent 7-Amino-4-Methyl Coumarin (AMC) released by DPP4 enzymatic activity in Caco2 cells treated with PBS (Control; n=4) or DMSO or DPP4 inhibitors (n=4). Results were analyzed using Kruskal-Wallis test with Dunn's post-test (**A**) from three independent experiments. *P<0.05; **P<0.01; ***P<0.001; ****P<0.0001. ns=not significant. Bars indicate mean of all data points.

**Flow cytometry gating strategy:** Cells were gated on FSC/SSC to select the entire population of cells, then gated on FSC-A/FSC-H to select for single cells. Flow plots shown reflect the entire cell population without subgating. To assess for HAstV infection, gating for positive vs. negative cells was done using uninfected vs. infected control cells and subsequently applied uniformly to experimental cells to avoid bias.
